# Left ventricular geometry during unloading and the end-systolic pressure volume relationship: Measurement with a modified real-time MRI-based method in normal sheep

**DOI:** 10.1101/562017

**Authors:** Duc M. Giao, Yan Wang, Renan Rojas, Kiyoaki Takaba, Anusha Badathala, Kimberly A. Spaulding, Gilbert Soon, Yue Zhang, Vicky Y. Wang, Henrik Haraldsson, Jing Liu, David Saloner, Liang Ge, Julius M. Guccione, Arthur W. Wallace, Mark B. Ratcliffe

## Abstract

The left ventricular (LV) end-systolic (ES) pressure volume relationship (ESPVR) is the cornerstone of systolic LV function analysis. We describe a 2D real-time (RT) MRI-based method (RTPVR) with separate software tools for 1) semi-automatic level set segmentation (LSSM) of the LV, 2) generation of synchronized pressure area loops and 3) calculation of the ESPVR. We used the RTPVR method to measure ventricular geometry, ESPAR and ESPVR during vena cava occlusion (VCO) in normal sheep.

10 adult sheep were anesthetized and 2D cross sectional RTMRI performed at apex, mid-ventricle and base levels during separate VCOs. The Dice similarity coefficient was used to compare LSSM and manual image segmentation and thus determine LSSM accuracy. LV cross-sectional area, major and minor axis length, axis ratio, major axis orientation angle and ESPAR were measured at each LV level. ESPVR was calculated with a modified Simpson’s rule method.

The Dice similarity coefficient between LSSM compared and manual segmentation by two readers was 87.31±2.51% and 88.13±3.43% respectively. All cross sections became more elliptical during VCO. The major axis orientation shifted during VCO but remained in the septo-lateral direction. LV chamber obliteration at the apical level occurred during VCO in 7 of 10 sheep. ESPAR was non-linear at all levels. Finally, ESPVR was non-linear because of apical collapse.

LSSM segmentation of 2D RT MRI images is accurate and allows calculation of LV geometry, ESPAR and ESPVR during VCO. In the future, RTPVR will facilitate determination of regional systolic material parameters underlying ESPVR.

## Introduction

The left ventricular (LV) end-systolic pressure (ES) volume relationship (ESPVR) is the cornerstone of LV systolic function analysis [1, 2]. For instance, the slope of the ESPVR (end-systolic elastance (E_ES_)), which is usually measured during a transient reduction in preload (vena cava occlusion (VCO)) [3], is widely used in basic and clinical research as a load independent measure of global LV systolic function [4].

However, there are several gaps in our knowledge of ESPVR. First, at the myocyte level, it is generally accepted that the active force myocyte/sarcomere length relationship determines the ESPVR [5, 6] but little is known about how underlying systolic material properties relate to ESPVR. Second, LV systolic function is regionally heterogeneous in both the normal LV [7] and in patients with ischemic [8] and valvular heart disease [9, 10] but little is known about how regional systolic function contributes to the ESPVR. Imaging of the LV during VCO is a necessary prerequisite for investigation in both of these areas.

Conventional ESPVR methods including sonomicrometry [11] and the conductance catheter [12] provide limited LV geometry data. The 3D location of sonomicrometry transducers can be determined with the array localization method [13], however, the number of transducers is limited, wall thickness is usually measured at a single point [7] and calculation of LV geometry must assume an ideal LV shape [10]. Dahl et al measured both ESPVR with a conductance catheter and strain and strain rate with 2D echocardiography during VCO in normal pigs but LV geometry was not reported [14]. Segmental LV volume data can be obtained from a conductance catheter [15] but this provides only indirect information about segmental LV shape.

Although conventional cine cardiac magnetic resonance imaging (CMRI) has excellent spatial resolution, acquisition times were not previously adequate for imaging during preload reduction. However, Zhang et al recently described a real-time CMRI (RTMRI) technique based on a radial FLASH sequence with nonlinear inverse reconstruction that has image acquisition times as short as 20 to 30 ms [16] and Witschey et al used the RTMRI method to acquire 2D images of the LV during VCO in normal sheep [17]. LV pressure area loops were created and the end-systolic pressure area relationship (ESPAR) was measured at a range of levels between the LV apex and base [18]. Nevertheless, the alterations in LV geometry during VCO have not been studied previously.

In the current study we describe a 2D RTMRI-based method (RTPVR) that includes separate software tools for 1) semi-automatic level set segmentation (LSSM) of the LV cavity, 2) generation of synchronized pressure area loops and 3) calculation of the ESPVR. We then use the method to measure ventricular geometry during VCO and corresponding ESPAR and ESPVR in normal sheep.

## Methods

### Overview

Software tools for 1) semi-automatic level set segmentation (LSSM) of the LV (Matlab R2016b, Mathworks, Natick, MA, 2) generation of synchronized pressure area loops and 3) calculation of the ESPVR were created (C#, Visual Studio 2017, Microsoft, Redmond, WA with Matlab.net compiler). We first validated the LSSM method of LV cavity segmentation by comparing manual segmentation performed by two image readers with LSSM output. The Dice similarity coefficient, which determines the similarity of two image objects as the ratio of twice the object intersection over the union of the image objects, is commonly used to determine the accuracy of image segmentation methods [19]. Accordingly, intra- and inter-observer variabilities and the Dice coefficient were calculated to determine the accuracy of the LSSM method.

A flowchart illustrating the RTPVR method is shown in **Fig 1.** Briefly, separate 2D short axis RT MRI was performed at apex, mid-ventricle and base LV levels during separate VCOs. LSSM calculated LV short axis geometry was used to calculate major and minor axes, axis ratio and axis orientation. LV pressure and area are then synchronized and ESPAR loops are generated. Mixed model regression was used to determine the effect of LV level on major and minor axes, axis orientation and ESPAR slope and intercept. ESPVR is then calculated using a modified Simpson’s rule method.

**Figure 1:**
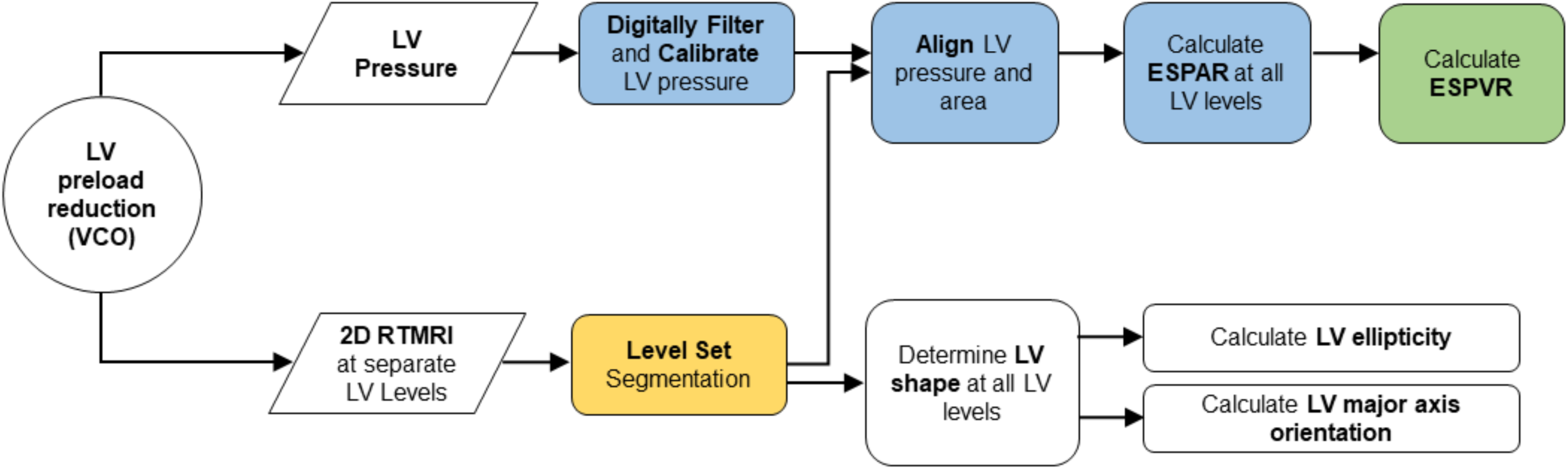
A flowchart illustrating the overall methodology designed to derive subject-specific ESPVR from RTMRI and LV pressure measurement. 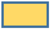-semi-automatic level set segmentation (LSSM) software module 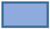-synchronized pressure area loop generation software module 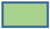-ESPVR calculation software module

### Cardiac MRI (CMRI)

Studies were approved by the San Francisco VA Institutional Animal Care and Use Committee (IACUC), in compliance with the “Guide for the Care and Use of Laboratory Animals” prepared by the Institute of Laboratory Animal Resources, National Research Council.

Ten healthy, adult sheep were sedated with ketamine (20mg/kg intravenous), anesthetized (Isoflurane 2.2% inhaled) and mechanically ventilated. The tip of a pressure catheter (SPC-350; Millar, Houston, TX) was immersed in water at 38°C for 12 hours prior to calibration and positioning in the LV and an 8 Fr balloon catheter was positioned in the inferior vena cava via femoral vessels using fluoroscopic guidance. Ferumoxytol (0.125 ml/kg IV over 1 hour; AMAG Pharmaceuticals, Waltham, MA) was given 1 hour prior to MRI [20]. Metoprolol (5 mg) and atropine (1 mg) were also given intravenously immediately prior to MRI. Isoflurane was maintained at 2.2%; end-tidal CO_2_ was kept between 25 and 45 mm Hg; an infusion of neosynephrine was titrated to keep peak LV pressure at 90+5 mm Hg during cardiac MRI.

MRI imaging was performed on a 3T MRI scanner (Skyra; Seimens, Verlagen, Germany). Six cine long axis MR images that were 30° apart were first obtained and this set of baseline scans was used for LVV determination prior to VCO [21]. 2D short axis RTMRI was performed 25 (Apex), 50 (Mid) and 75% (Base) of the distance from the apex to the base of the LV during separate VCOs. The MR imaging parameters are summarized as follows: 2D multislice retrospectively-gated cine balanced steady-state free-precession acquisition with the following imaging parameters, TE = 1.34 ms, TR = 59.2 ms, acquisition matrix = 128 × 54, FOV = 178 × 260, slice thickness = 8 mm, pixel spacing = 2.0313 × 2.0313 mm.

LV pressure (LVP) acquisition (12 bit, 5K samples/ sec, ACQ16 and Ponemah 5.2, DSI, St. Paul, MN) started approximately 5 seconds before the start of the MRI acquisition as shown in **Fig 2**. LV pressure was filtered with an in-line analog filter (BNC Low Pass Filter, Crystek, Fort Myers, FL) during acquisition and post-processed with a digital constrained least Pth-norm IIR filter (Matlab). LVP during VCO was calibrated using pressure acquired immediately prior to MRI acquisition as shown in **Fig 2B**.

**Figure 2:**
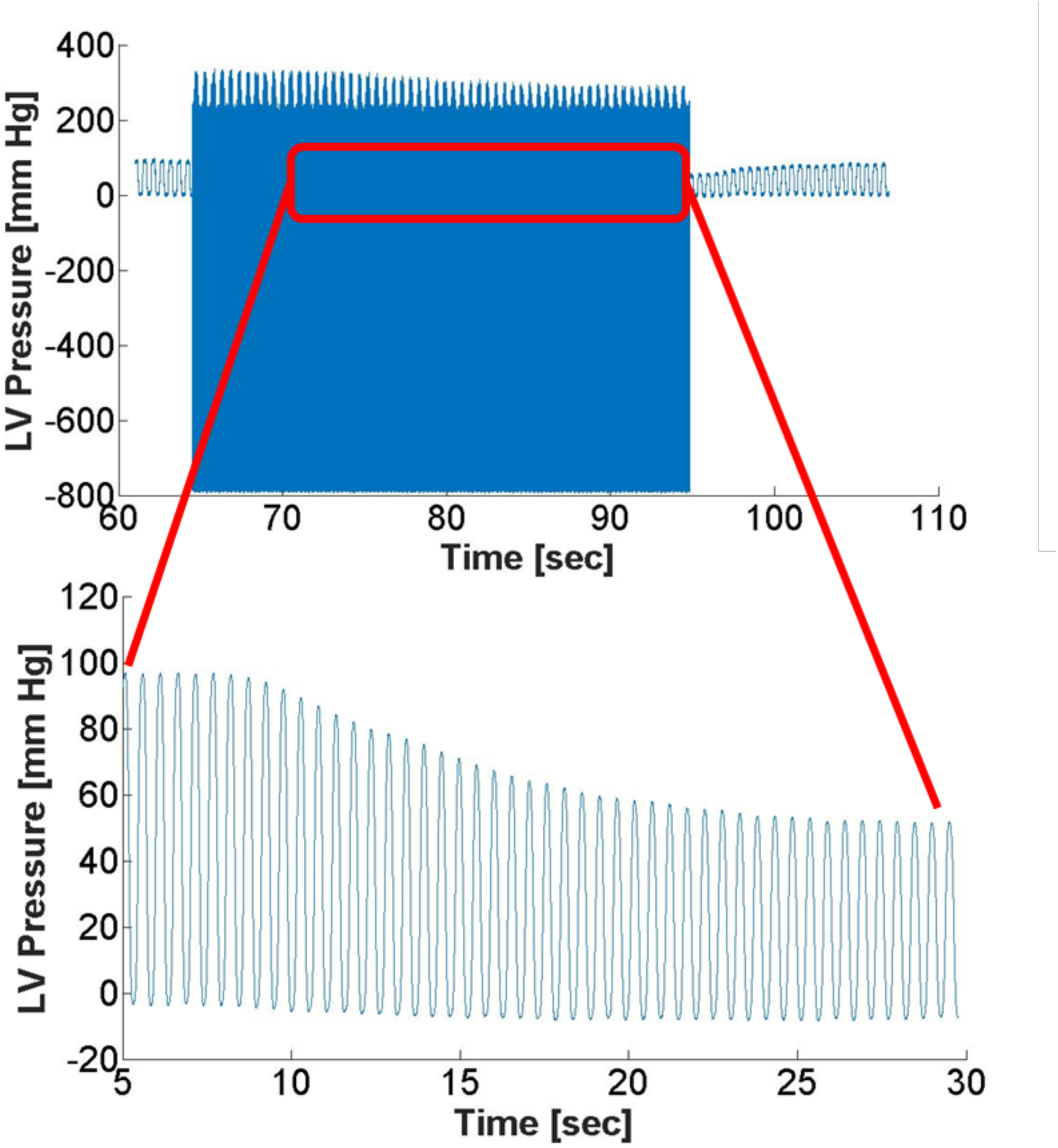
Raw pressure data with noise prior to digital filtering and calibration (top) with the insert showing filtered and calibrated LV pressure during the selected VCO range.

### Validation of semi-automatic level-set segmentation

Manual segmentation was used as the ground-truth to validate the accuracy of the level set algorithm [22, 23]. Manual segmentation was performed in a blinded manner. Two readers drew manual contours for 200 slices from all 10 subjects (20 slices for each case) as shown in **Fig 3A**. A second set of manual contours was drawn by each of the readers in 3 of the cases allowing the calculation of both inter- and intra-observer variability. The Dice similarity coefficient, commonly used to determine the accuracy of image segmentation methods, was used to determine the accuracy of the LSSM contours (**Fig 3B**) [19].

**Figure 3:**
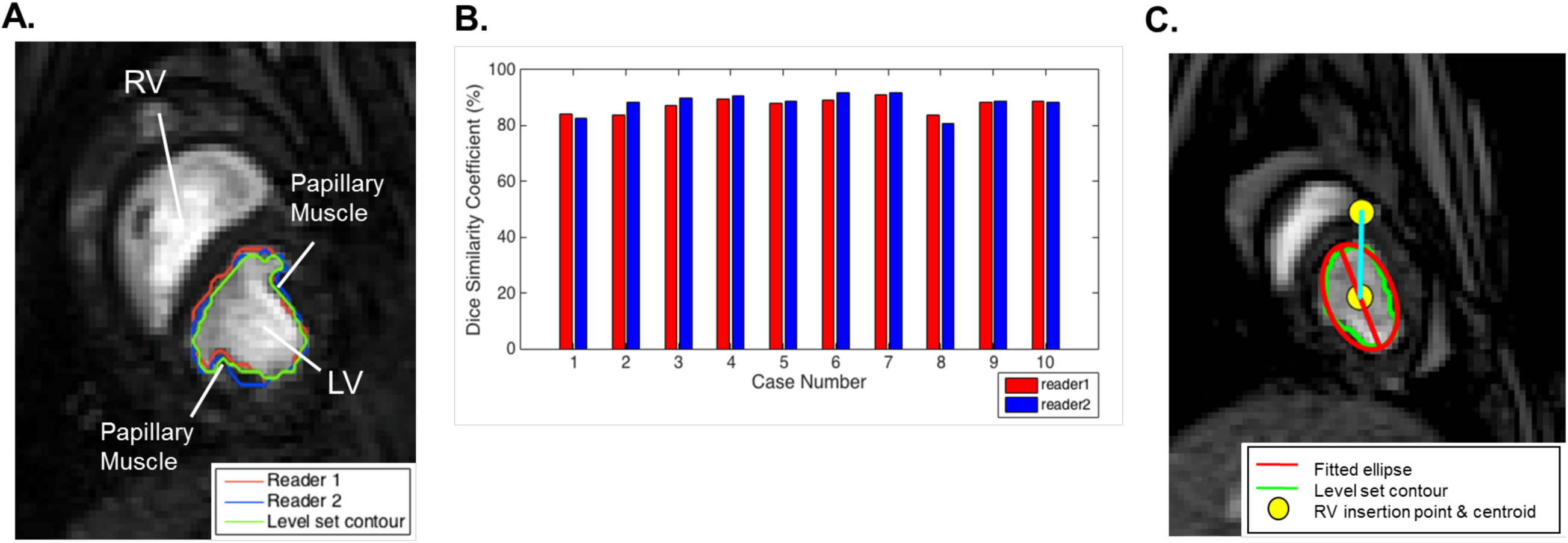
**(A)** Representative midventricular short-axis image of the LV showing level set and manual contours. Green = automatic contouring, Red = manual contour reader 1, Blue = manual contour reader 2. **(B)** Dice similarity coefficient for all 10 cases. (**C**) Representative midventricular short-axis image of the LV overlaid with automatic contours and fitted ellipse for quantifying major/minor axis length and major axis orientation angle as surrogate measurement of LV shape change. Red = fitted ellipse. Green = automatic contouring. Yellow circle = RV insertion point and centroid of ellipse. Movie is included in the Supplementary Material.

### Image segmentation

The LSSM was used to segment the 2D RTMRI images [24]. Specifically, the first frame of each imaging series was manually contoured while subsequent frames were processed with the LSSM [24].

LSSM generated contours were used to calculated LV cross-sectional area. In addition, an ellipse was fit to the LSSM generated contours and major and minor axes, axis ratio, and major axis orientation angle were calculated. Note that the orientation was relative to the anterior right ventricular endocardial insertion and the LV septum was in the positive direction **(Fig 3C)**.

Obliteration of the LV cavity was defined as LV area ≤ 0.25 cm^2^. In instances where the LV cavity collapsed, axis and angle data were excluded. Representative LV shape and area changes during VCO are shown in **Fig 4.**

**Figure 4:**
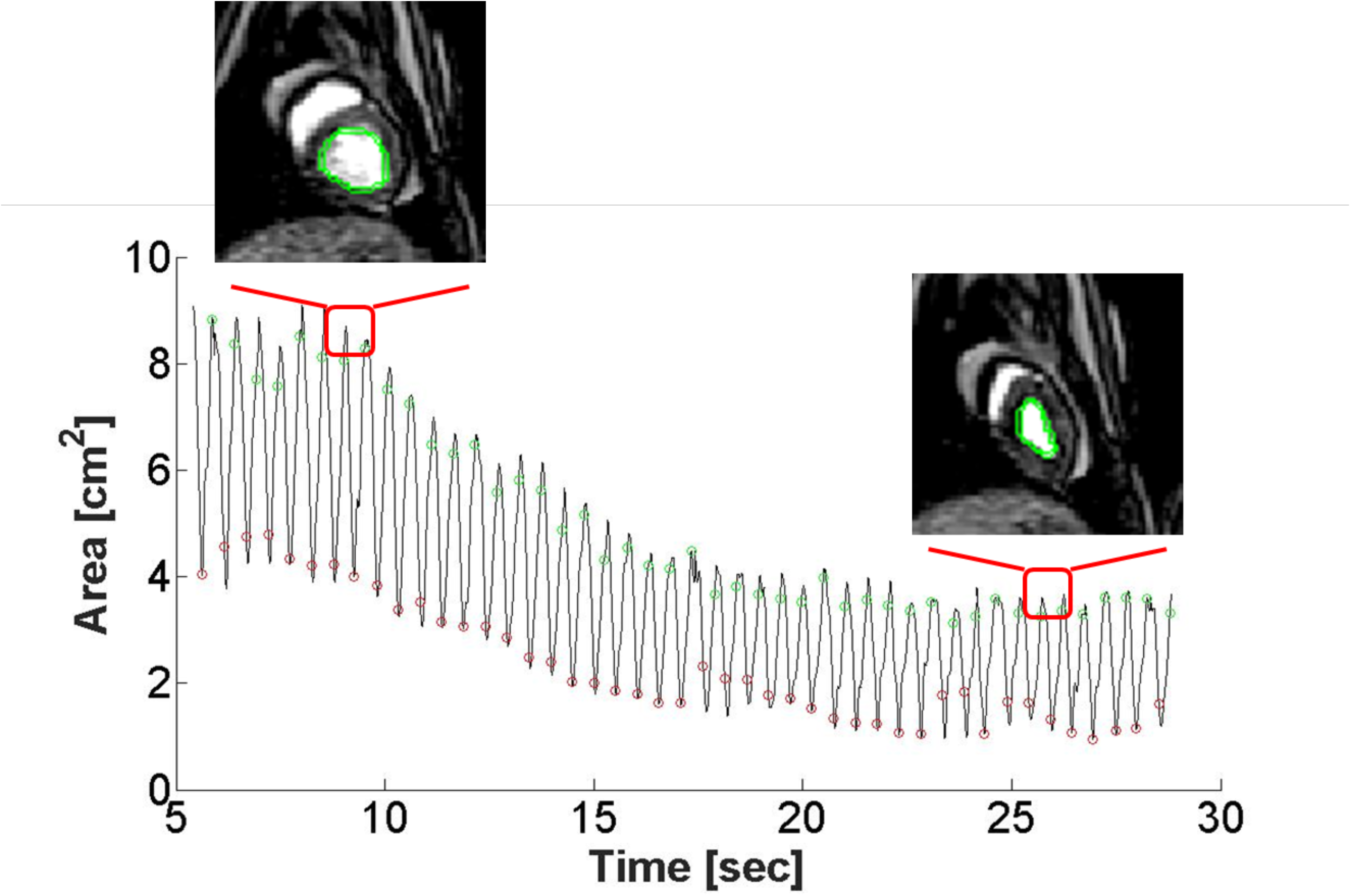
LV Area plot calculated by the level set algorithm with representative midventricular short-axis images of the LV before VCO and during VCO starting at t = 10 sec. Green makers = end-diastole (ED), Red markers = end-systole (ES).

### ESPAR calculation

LV area was initially aligned to the region of noise on the unfiltered LVP trace. The time derivative of the LVP (d LVP/dt) was calculated and 0.25 peak d LVP/dt was aligned with peak LV area. The alignment method was chosen to obtain optimal LV PA loops (**S1 Figure**). The ESPAR was calculated using a shooting point iterative regression approach previously described by Kono et al for ESPVR determination from conductance catheter data [25].

### ESPVR calculation

LV volume at ES was determined with the following:

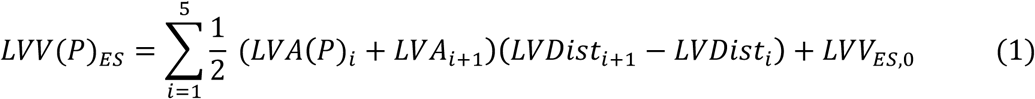

where *LVA*(*P*)_*i*_ is LV cross-sectional area with i ranges from 1 (LV apex) to 5 (LV valve plane), *LVDist*_*i*_ is the distance of the real time MRI short-axis slices from the LV apex. Note That *LVDist*_5=*ValvePlane*_ is assumed to move toward the apex in line with sonomicrometry data obtained in conscious dogs during VCO [26]. *LVV*_*ES,0*_ is the baseline ES volume calculated from step 1 and is assumed to be constant during the VCO. Subsequent points on the ESPVR are generated using *LVA*(*P*)_*i*_ at LVP range between 50 and 90 mmHg with a 5 mm Hg step change.

ESPAR and ESPVR calculation was performed with custom code written in C# (Visual Studio 2017, Microsoft, Redmond, WA) using Matlab routines (.NET assembly using Matlab Compiler, Mathworks).

### Statistical analysis

All values are expressed as mean ± standard error of the mean. The significance level was set at p<0.05. Multivariate mixed effect analyses (Proc Mixed, SAS version 9.2, SAS Institute Inc., Cary, NC) were used to determine variable change during VCO. Individual sheep were included as a random effect [27]

## Results

### Semi-automatic level set method

LSSM-based segmentation was automatic in 7 of 10 sheep data sets and required minor parameter modifications in the remaining 3 of 10 data sets. The intra-observer variability from the two readers were 0.54% and 0.88%. The Dice similarity coefficient was used for inter-observer variability measurement between level set segmentation comparing with manual results from two readers and determined to be 87.31%±2.51% and 88.13%±3.43% respectively, thus showing achieving a high segmentation accuracy as shown in **Fig 3B**. A representative LV area vs time plot is shown in **Fig 4**.

### Hemodynamic parameters

Cycle length, ES interval and LV pressure at baseline are seen in **Table 1**. The ES interval was significantly shorter at the mid ventricle level (0.246+0.005 sec) than at the apical level (0.261+0.005 sec) but the cardiac cycle length was not different between LV levels. At baseline, the LVP was significantly lower at the apical level (88.6 ± 1.60 mm Hg) than both the mid-ventricle and basal levels (94.8 ± 1.34 and 95.1 ± 1.40 mm Hg respectively).

**Table 1.** Baseline Cardiac MRI Measurements.

|  | **All VCOs**<br>**(n=30)** | **Apex**<br>**(n=10)** | **Mid**<br>**(n=10)** | **Base**<br>**(n=10)** | **Significance\*** |
| --- | --- | --- | --- | --- | --- |
| Cycle Length [sec] | 0.637+0.007 | 0.631 + 0.008 | 0.640 + 0.012 | 0.641+ 0.014 | NS |
| ES Interval [sec] | 0.255+0.003 | 0.261 + 0.005 | 0.246 + 0.005 | 0.257 + 0.005 | 1, 3 |
| LV Pressure at ES [mm Hg] | 92.9+0.9 | 88.6 + 1.6 | 95.1 + 1.4 | 94.8 + 1.3 | 2, 3 |
| LV Area at ES [cm<sup>2</sup>] | 5.43+0.25 | 2.53 + 0.156 | 4.55 + 0.181 | 9.15 + 0.323 | 1, 2, 3 |
| LV Major Axis at ES [cm] | 2.77+0.06 | 1.99 + 0.0601 | 2.637 + 0.0573 | 3.66 + 0.0627 | 1, 2, 3 |
| LV Minor Axis at ES [cm] | 2.37+0.06 | 1.63 + 0.0622 | 2.27 + 0.0456 | 3.20 + 0.0576 | 1, 2, 3 |
| LV Axis Ratio at ES | 1.19+0.01 | 1.26 + 0.0307 | 1.16 + 0.0150 | 1.15 + 0.0129 | 2, 3 |
| Major Axis Orientation Angle at ES [°] | 31.7+3.7 | 13.7 + 7.46 | 51.2 + 5.67 | 29.9 + 4.74 | 1, 2, 3 |

Baseline cycle length, end-systolic (ES) interval, LV pressure and area, major and minor axis, axis ratio and axis angle at ES for all VCO data and at three LV levels. \*NS: not significant, 1: Base vs Mid (p<0.05), 2: Base vs Apex (p<0.05), 3: Mid vs Apex (p<0.05).

Changes in cycle length, ES interval and LV pressure during VCO are seen in **Table 2.** There was statistically significant shortening of both the ES interval (−0.0038+0.0005 sec/ beat) and cardiac cycle length (−0.0024+0.0004 sec/ beat) during VCO. For reference, this is a 4.8 msec decrease in cycle length over a 20 beat VCO. The rate of change of LVP during VCO was not different between LV levels.

**Table 2.** VCO Cardiac MRI Measurements.

|  | **All VCOs**<br>**(n=30)** | **Apex**<br>**(n=10)** | **Mid**<br>**(n=10)** | **Base**<br>**(n=10)** | **Significance\*** |
| --- | --- | --- | --- | --- | --- |
| Cycle Length [sec/ beat] | -0.0038+0.0005 | -0.0027+0.0008 | -0.0043+0.0011 | -0.0042+0.0006 | 1 |
| ES Interval [sec/ beat] | -0.0024+0.0004 | -0.0035+0.0009 | -0.0018+0.0008 | -0.0020+0.0004 | 1 |
| LV Pressure at ES [mm<br>Hg/ beat] | -2.26+0.11 | -2.32+0.19 | -2.23+0.21 | -2.22+0.18 | 1 |
| LV Area at ES [cm<sup>2</sup>/ beat] | -0.13+0.02 | -0.08+0.02 | -0.11+0.01 | -0.21+0.04 | 1,2 |
| LV Major Axis at ES [cm/<br>beat] | -0.038+0.003 | -0.033+0.008 | -0.040+0.004 | -0.043+0.0058 | 1 |
| LV Minor Axis at ES [cm/<br>beat] | -0.042+0.004 | -0.033+0.009 | -0.041+0.004 | -0.050+0.008 | 1 |
| LV Axis Ratio at ES | 0.0080+0.0017 | 0.0084+0.0041 | 0.0072+0.0020 | 0.0085+0.0026 | 1 |
| Major Axis Orientation<br>Angle at ES [°/ beat] | -0.19+0.28 | -1.04+0.46 | -0.34+0.54 | 0.81+0.27 | 1,2 |

Slope of cycle length, end-systolic (ES) interval, LV pressure and area, and major and minor axis, axis ration and axis angle at ES vs
VCO beat for all VCO data and at three LV levels. \*1: All VCO variables vs beat (p<0.05), 2: Base vs Apex (p<0.05).

### LV geometry

LV cross-sectional area, major and minor axes, axis ratio, and major axis orientation are shown in **Table 1**. As expected, LV area, major axis, and minor axis lengths increased from the apical to the basal segment. Conversely, axis ratios decreased from the apical to the basal segment, confirming that the LV is more spherical at the basal segment and more elliptical at the apical segment. Major axis orientation angle was greatest at the mid-ventricle level with an average angle of 51.2° while the base and apex levels had average angles of 29.9° and 13.7°respectively.

The change in LV cross-sectional area, major and minor axes, axis ratio, and major axis orientation during VCO are shown in **Table 2**. Also, **Fig 5** shows representative changes in major axis, minor axis, axis ratio, and major axis orientation angle at mid-ventricle LV level during VCO. Briefly, all 5 parameters changed significantly during VCO. The significant change in axis ratio documents that the LV becomes more elliptical during VCO and the significant change in major axis orientation shows that the major axis shifts during VCO but remains generally oriented in the septo-lateral direction.

**Figure 5:**
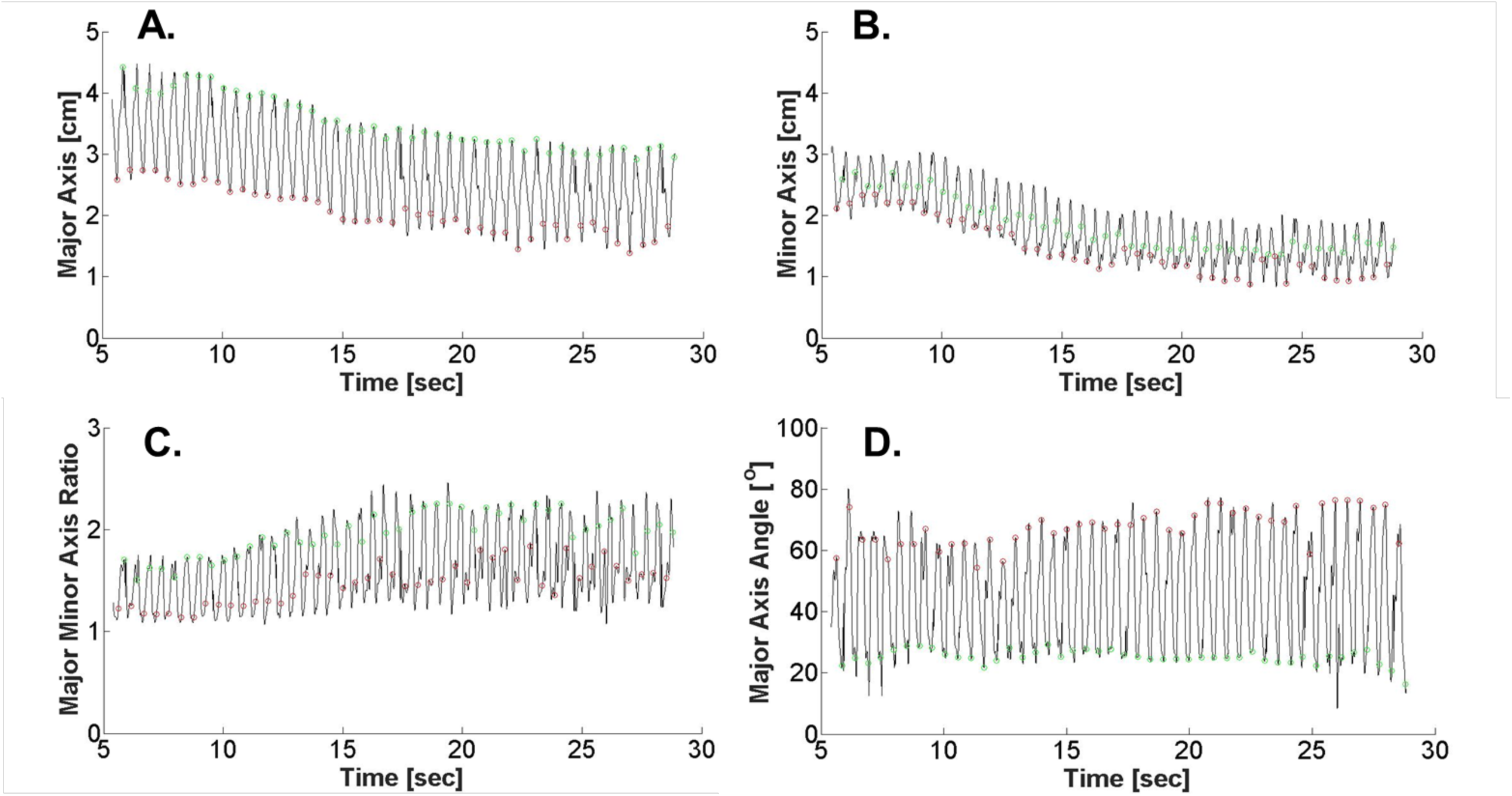
Changes in LV geometry before and during VCO. Shape changes were quantified by 4 indicators calculated from the fitted ellipse:**(A)** Major Axis length, **(B)** Minor Axis length, **(C)** Major to Minor Axis Ratio (MajMinR), and **(D)** Major Axis Orientation Angle. Green makers = end-diastole (ED), Red markers = end-systole (ES).

There were statistically significant differences between LV levels in the rate of change of LV area (−0.08+0.02 cm^2^/ beat apex vs -0.21+0.04 cm^2^/ beat base level) and major axis orientation (−1.04+0.46°/ beat apex vs 0.81+0.27° / beat base) during VCO. There was no difference in major and minor axis or axis ratio between LV levels during VCO.

Last, LV collapse at the apical segment during VCO was observed in 7 of the 10 cases (27 beats). **Fig 6** shows representative plots of LV area, major axis, and minor axis at the apical level. In each case, the dashed red line represents the time when LV cross sectional area fell below 0.25 cm^2^.

**Figure 6:**
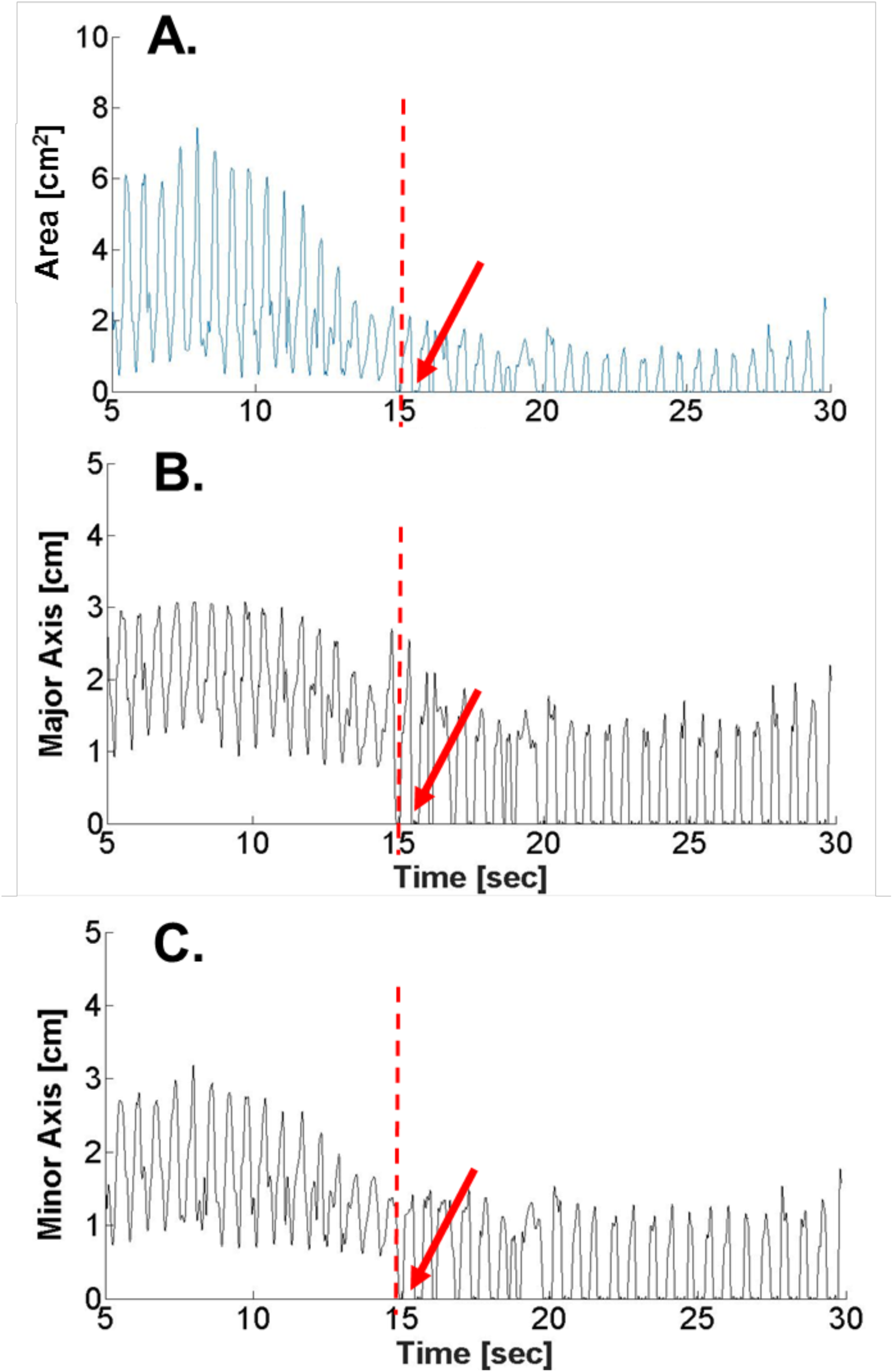
Plots of LV Apical Area **(A)**, Major Axis length **(B)**, and Minor Axis length **(C)** of a case showing segmental collapse where LVA < 0.25 cm^2^ after time = 15 sec as indicated by a red arrow. The vertical dashed line represents collapse of the LV to zero volume.

### ESPAR

Fig 7 shows representative LV pressure area loops during VCO at the apex, mid-ventricle and base levels. PA loops shift to the left and the slope of the ESPAR becomes more steep as the LV level moves from base to apex.

ESPAR slope and LV pressure intercept are illustrated in **Fig 8A, Fig 8B**, and **S1 Table**. The LV pressure intercept for the apex level was significantly higher than mid-ventricle (12.02 mm Hg) and base (8.39 mm Hg) levels mirroring the fact that the apical cavity obliterates before mid and base levels. The mixed-effect model showed that the ESPAR was non-linear (β_2_= -0.500 (p<0.001), β_1_=9.209 (p<0.001), β_0_=48.73 (p<0.001)).

**Figure 7:**
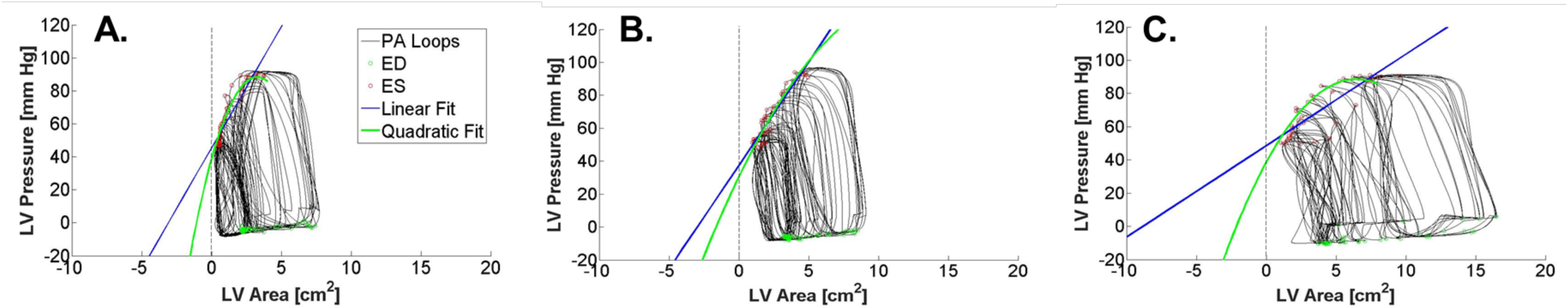
Representative LVP-area (LVPA) loops derived from the level set algorithm and pressure measurements along with fitted ESPAR at apex **(A)**, mid-ventricle **(B)**, and base **(C)** during VCO. The ESPAR was fitted with a linear function (blue) and a quadratic function (green) for comparison. The vertical dashed line represents collapse of the LV to zero volume.

**Figure 8:**
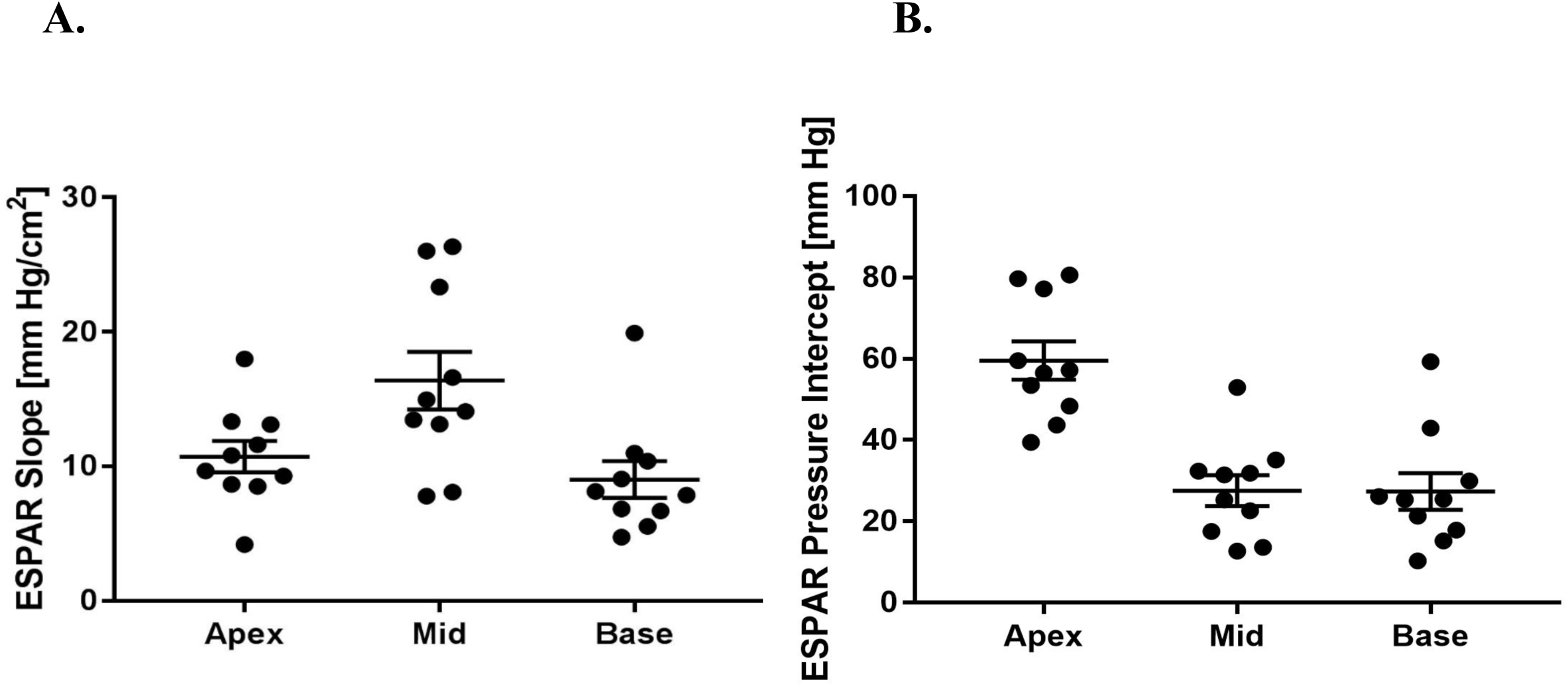
Slope of ESPAR **(A)** and LV area intercept A_o_ **(B)** at apex, mid, and base LV levels for all animals indexed to basal surface volume.

### ESPVR

ESPVR, calculated using modified Simpson’s rule (**Equation 1**), are shown in **Fig 9** and **S2 Table.** A representative ESPVR is shown in **Fig 9A.** Overall, we found E_ES_ to be extremely reproducible between animals (E_ES_ = 2.31 ± 3.1 mm Hg/ml **(9B)**; V_o_= -13.6 ±2.41 mL **(9C)**). The mixed-effect model showed that the ESPVR is nonlinear (β_2_=0.0414 (p<0.001), β_1_=0.349 (p<0.001), β_0_=18.12 (p<0.001)).

**S1 Table.** Slope and volume intercept of ESPAR.

|  | **ESPAR Slope [mm Hg/cm<sup>2</sup>]** |  |  | **A<sub>o</sub> [cm<sup>2</sup>]** |  |  |
| --- | --- | --- | --- | --- | --- | --- |
| **Animal #** | **Apex** | **Mid** | **Base** | **Apex** | **Mid** | **Base** |
| 1 | 4.18 | 26.3 | 11.0 | -18.5 | -0.858 | -1.63 |
| 2 | 11.6 | 23.4 | 9.07 | -4.87 | -0.543 | -2.80 |
| 3 | 9.67 | 8.08 | 6.68 | -4.52 | -0.395 | -2.28 |
| 4 | 18.0 | 16.6 | 8.13 | -2.97 | -0.820 | -3.69 |
| 5 | 13.3 | 13.5 | 4.75 | -3.63 | -2.61 | -12.5 |
| 6 | 9.28 | 7.80 | 6.84 | -4.25 | -4.04 | -1.50 |
| 7 | 10.8 | 26.0 | 19.9 | -5.29 | -0.675 | -1.07 |
| 8 | 8.67 | 13.1 | 7.86 | -9.20 | -4.03 | -3.32 |
| 9 | 13.1 | 15.0 | 5.54 | -4.55 | -1.69 | -7.75 |
| 10 | 8.51 | 14.1 | 10.4 | -9.48 | -2.30 | -2.46 |
| Mean + SEM | 10.7 + 1.16 | 16.4 + 2.13 | 9.02 + 1.36 | -6.73 + 1.48 | -2.15 + 4.59 | -3.90 + 1.12 |

**S2 Table.** End-systolic elastance E_ES_ and volume-axis intercept.

| **Animal #** | **$E_{ES}$ [mm Hg/mL]** | **$V_o$ [mL]** |
| --- | --- | --- |
| 1 | 3.26 | -8.21 |
| 2 | 1.87 | -16.0 |
| 3 | 1.38 | -18.4 |
| 4 | 2.35 | -12.7 |
| 5 | 1.74 | -29.2 |
| 6 | 1.57 | -15.4 |
| 7 | 2.51 | -11.2 |
| 8 | 1.96 | -16.6 |
| 9 | 1.79 | -6.64 |
| 10 | 4.70 | -7.73 |
| Mean + SEM | 2.31+3.10 | -13.6+2.41 |

**Fig 9:**
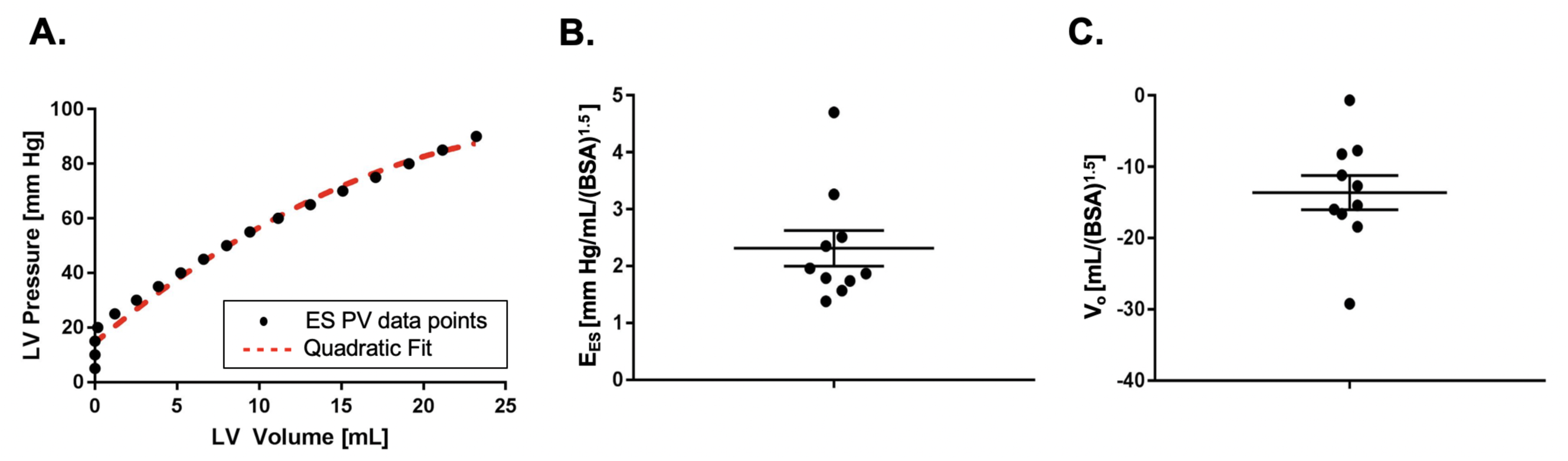
Representative ESPVR with a quadratic fit in red dashed line **(A).** End-systolic elastance E_ES_ **(B)** and volume intercept V_o_ **(C)** of ESPVR indexed to basal surface volume.

## Discussion

The principal finding of the study is that the RTPVR method allows measurement of LV geometry during VCO and calculation of corresponding ESPAR and ESPVR in normal sheep. Specific findings included that the LV shape becomes more elliptical during VCO leading to eventual apical LV cavity obliteration and that ESPAR and ESPVR are non-linear.

### Semi-automatic active contouring

In the current study, an LSSM software tool was used to semi-automatically segment the LV during VCO. Briefly, semi-automatic LSSM generated contours were comparable to manual segmentation. The computation time for semi-automatic segmentation was less than 10 seconds per slice, although more powerful computing could greatly shorten the processing time.

### LV geometry and apical level cavity obliteration during VCO

The effect of VCO on LV shape during ESPVR measurement has not been previously studied. However, similar to our findings, existing LV geometry data changes in a non-concentric fashion during VCO. For instance, Olsen et al. showed that the anterior posterior (AP) minor axis decreases more than the septal-lateral (SL) minor axis during VCO [28]. Furthermore, LV cavity obliteration occurs in the setting of intra-operative hypovolemia [29] and it is therefore seemed likely that LV cavity obliteration occurs during VCO.

The reason for the increased LV ellipticity during VCO where the major axis orientation is in the septo-lateral direction is unclear but could be due to either differential structural stiffness or transmural pressure in the interventricular septum. We did not measure right ventricular (RV) pressure. However, Olsen et al measured LV and RV pressure during VCO in conscious normal dogs and found that the ratio of LV to transmural septal pressure at ES is relatively constant (Calculated from Olsen Figure 5) [30]. Further work is warranted but this finding suggests that the septum may be structurally different than the LV free wall.

Contijoch et al. observed a similar increase in ESPAR pressure intercept at the LV apex in their real time MRI based study in normal sheep. However, that study did not observe cavity obliteration at the LV apex [31]. We used a neosynephrine infusion to counteract anesthesia related hypotension and maintain peak LV pressure at 90 mm Hg. As a consequence, LV pressure change during VCO in our study was somewhat higher. This may have been a factor and it is also possible that the volume status of sheep in the two studies was different.

### ESPVR

Implicit in ESPVR analysis is the assumption that there is a relationship between the pressure volume relationship and regional systolic myocardial parameters [1]. Determination of that relationship would be complex and would probably require an inverse finite element (FE) solution where LV geometry and cavitary pressure are input and regional contractility is adjusted so that calculated myocardial deformation agrees with measured values [32]. However, such a FE based determination of the ESPVR/ regional contractility relationship would not be possible without imaging-based determination of LV geometry during VCO.

It is now generally accepted that the ESPVR is non-linear. For instance, van der Velde found that ESPVR is convex with center of curvature to the right and quadratic term (β_2_) approximately -0.3 mm Hg/ ml^2^ in normal pigs (Calculated for control values from van der Velde Table 2) [33]. We found that ESPAR at all LV levels is significantly non-linear. We also found that ESPVR is significantly non-linear with curvature to the right but with β_2_ = -0.0414. Further study is warranted but non-linearity of ESPAR and the difference in β_2_ between our study and van der Velde’s suggests that ESPVR non-linearity is the product of both underlying material properties and apical cavity obliteration.

### Limitations

With the current method we are able to acquire 6-8 images per beat which we interpolate to create PA loops. Further, synchronization between pressure and area is based on pressure and area metrics rather than the ECG.

Last, LSSM segmentation of the LV long axis during VCO was thought to not be possible because of algorithm failure to distinguish LV and left atrium at the mitral valve level.

### Conclusions and future directions

In conclusion, LSSM segmentation of 2D RT MRI images is accurate and allows measurement of LV geometry during VCO and calculation of corresponding ESPAR and ESPVR. Although the goal was to describe the RTPVR method, it was noted that LV shape becomes more elliptical during VCO leading to eventual apical LV cavity obliteration and that ESPAR and ESPVR are non-linear.

Future work will focus on semi-automatic segmentation of the LV long axis during VCO with machine learning based methods. Hopefully, more rapid image acquisition as well as actual 3D RTPVR methods will also be developed. Last, similar RTPVR methods should be developed for the end-diastolic pressure volume relationship and, as before, the RTPVR will facilitate determination of the relationship between ESPVR and underlying regional systolic material parameters.

## Acknowledgements

The authors would like to acknowledge the staff from the animal facility at the San Francisco Veterans Affairs Medical Center for their help facilitating the animal procedures. This study was supported by National Heart, Lung and Blood Institute Grant R01-HL-084431 (M. Ratcliffe).

---

**S1 Figure:**
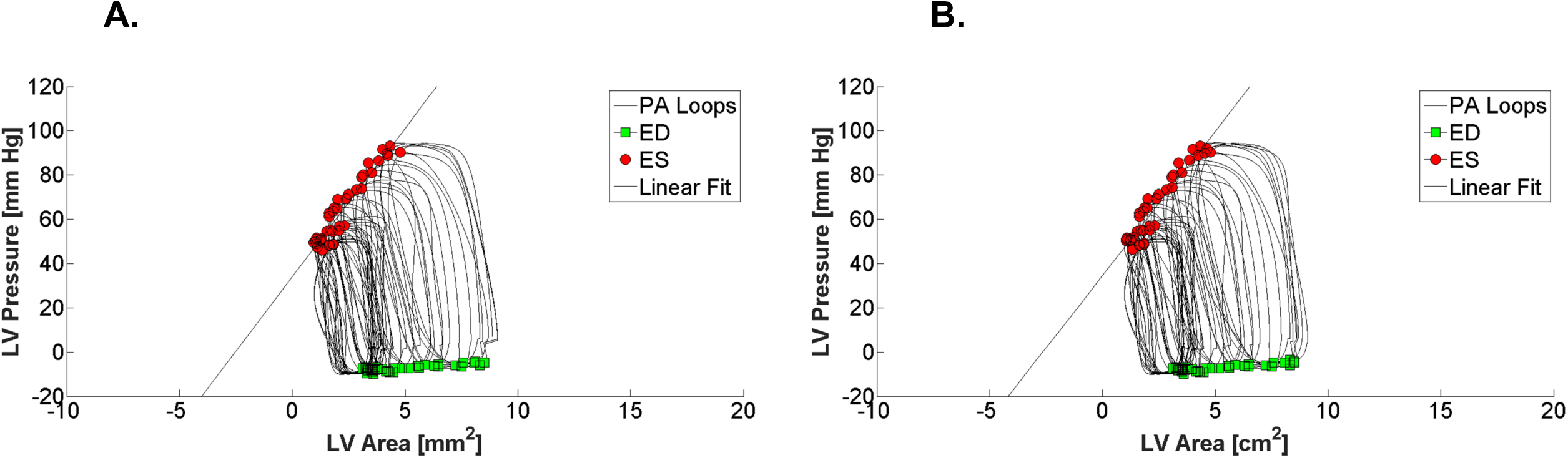
Alignment using max + dLVP/dt and max LV area was initially chosen because of the perception that beat shape was more appropriate. In retrospect, there is little effect between this (A) and 0.25 max + dLVP/dt and max LV area (B).

## Supporting Information

**S1 Figure:** Alignment using max + dLVP/dt and max LV area was initially chosen because of the perception that beat shape was more appropriate. In retrospect, there is little effect between this (A) and 0.25 max + dLVP/dt and max LV area (B).

**S1 Table**: Slope and volume intercept of ESPAR at three LV levels derived from RTMRI-based analysis.

**S2 Table**: End-systolic elastance E_ES_ (slope) and volume-axis intercept (V_o_) estimated from the ESPVR indexed to basal surface volume (BSA)^1.5^.

